# *Enterococcus faecium* colonization and persistence in a model of diabetic wound infection

**DOI:** 10.1101/2025.11.20.689639

**Authors:** Navin Jeyabalan, Frederick Reinhart Tanoto, Haris Antypas, Cheryl Neo Jia Yi, Rachel Tan Jing Wen, Kevin Pethe, David L. Becker, Claudia J. Stocks, Kimberly A. Kline

## Abstract

Chronic wound infections are a common comorbidity of diabetes mellitus and can progress to amputation if untreated, yet effective strategies to manage these infections are limited. Commensals such as *Enterococcus faecium* and *Staphylococcus epidermidis* can transition into opportunistic pathogens when host defenses are compromised, underscoring the complexity of chronic wound microbiology. *E. faecium*, particularly vancomycin-resistant strains, are an understudied, clinically important cause of chronic diabetic wound infections. Using a low-dose streptozocin-induced diabetic mouse model, we characterized *E. faecium* wound infection dynamics and identified differences in colonization and clearance compared to non-diabetic animals. At eight hours post infection (hpi), control mice exhibited higher *E. faecium* wound colony forming units (CFU) than diabetic mice, but cleared the infection more efficiently, resulting in similar CFU by 24 hpi. By contrast, diabetic mice showed impaired clearance, with elevated CFU persisting through 72 hpi. In mixed species infection with *S. epidermidis, S. epidermidis* CFU increased at 72 hpi while *E. faecium* CFU remained comparable to single species infection. Despite strong initial cytokine and neutrophil responses, *E. faecium* persisted in all wounds. Sustained neutrophil recruitment at 72 hpi occurred only in diabetic mice, whereas macrophage accumulation increased from 24 to 72 hpi in all wounds, including sterile controls. Histological analysis showed epithelial hyper-thickening in both groups, indicating that diabetes and *E. faecium* each contribute to impaired wound healing. This study establishes a diabetic mouse model of *E. faecium* wound infection and suggests that *E. faecium* modulates innate immune responses to persist in the wound bed.

## Introduction

Diabetes is a chronic metabolic disorder characterized by hyperglycemia resulting from partial or complete insulin insufficiency [1]. The burden of diabetes in adults is increasing: the global number of adults with diabetes rose to about 828 million in 2022 from an estimated 198 million in 1990 [2]. Individuals with diabetes are more susceptible to a range of infections including urinary tract infections (UTI), bacteremia, soft tissue infections, and cutaneous ulcerations, which if left untreated may progress to chronic wounds marked by persistent inflammation [3, 4]. Chronic wound infections are defined as non-healing wounds that show no significant reduction in size after two to four weeks of medical care [5]. These wound infections are often polymicrobial and pose a significant barrier to successful healing [5, 6]. Reduced pain perception in individuals with diabetes increases the risk of neglecting minor cuts and abrasions [7], increasing the likelihood of infection and progression to chronic wound states. Of particular concern are diabetic foot ulcers (DFU), which are difficult to treat, prone to infection, and frequently result in limb amputations [8]. A population-based study in the UK reported that 40% of individuals with diabetes from peripheral vascular disease developed foot ulcerations [9]. In Singapore, a qualitative study described the cost of managing diabetes-related amputation as up to $30,000 USD per patient [10].

DFU are frequently dominated by opportunistic bacteria including *Staphylococcus aureus*, *Pseudomonas aeruginosa*, *Streptococcus* spp. and Enterococci such as *E. faecalis* and *E. faecium* [11–13]. Among these human-infection associated Enterococci, most studies to date have focused on *E. faecalis*, however the increasing global prevalence of antibiotic resistant *E. faecium* demands urgent attention [14]. Its clinical importance is reflected in its inclusion in the ESKAPE group, which describes six multidrug-resistant bacterial pathogens that are major threats to hospitalized individuals [15]. *E. faecium* is also listed as a high-priority pathogen by both the WHO and CDC due to its resistance to last-line antibiotics and its role in healthcare-associated infections such as bacteremia and chronic wound infections [16, 17]. In addition to its intrinsic resistance to last-line antibiotics such as vancomycin, daptomycin and linezolid [18], clinical isolates of *E. faecium* often encode virulence factors associated with biofilm formation [19], which can further complicate the treatment of enterococcal wound infections [20, 21]. In a nursing facility, rectal swabs collected at the baseline visit revealed that 40% of patients were colonized with vancomycin-resistant *Enterococcus* species. Among these, 17.8% were colonized with vancomycin-resistant *E. faecium* and 8.4% with vancomycin-resistant *E. faecalis*, with some patients harboring both. Colonization with vancomycin-resistant *E. faecium* persisted for approximately twice as long as *E. faecalis* [22]. However, experimental models to study *E. faecium* infection dynamics remain scarce.

In this study, we characterized the infection dynamics and host response of vancomycin-resistant *E. faecium* strain E745 wound infection in a streptozocin-induced diabetic mouse model. *E. faecium* successfully colonized wounds in both control and diabetic mice and persisted for up to five days. While initial replication was greater in control mice, clearance was also more efficient. By contrast, *E. faecium* replication was not as apparent in diabetic wounds, but the clearance was not as effective, resulting in higher *E. faecium* numbers in diabetic wounds at 72 hpi. Diabetic wounds showed elevated neutrophil recruitment and impaired resolution of inflammation, whereas macrophage recruitment was comparable across groups. Despite early inflammatory cytokine induction, inflammatory signaling waned by 72 hpi despite the persistent presence of bacteria. Histological analysis revealed epithelial hyper-thickening, indicative of poor wound healing in *E. faecium* infected diabetic mice at 72 hpi. These findings suggest a role for *E. faecium* in delaying wound healing in diabetic wound infections.

## Results

### *E. faecium* persists in diabetic wounds

To model *E. faecium* diabetic wound infection, C57BL/6J mice were rendered hyperglycemic with streptozocin (STZ) [23]. All control mice remained below the diabetic threshold of 300 mg/dl of blood glucose, whereas STZ-treated mice were diabetic by day 15 (**Fig. S1A-B**). Full thickness, dorsal excisional wounds were inoculated with 10^6^ vancomycin-resistant *E. faecium* strain E745 and wound colony forming units (CFU) were quantified at defined timepoints. At 8 hours post infection (hpi), CFU in control wounds rose to ∼10^7^, indicating acute replication, while CFU in diabetic wounds were equivalent to inoculum (**Fig. 1A**). By 24 hpi, wound CFU in both groups declined to ∼5 x 10^5^ CFU (**Fig. 1B**). At 72 hpi, CFU in the control wounds dropped further to ∼10^3^, whereas CFU in the diabetic wounds stabilized at ∼10^4^ (**Fig. 1C**). Thus, bacterial numbers decreased gradually in control wounds from 8 to 72 hpi, but only declined between 8 and 24 hpi in diabetic mice, remaining stable thereafter (**Fig. 1D**). In preliminary experiments extended to 5 days post infection, wound CFU were ∼10^3^ in both groups (**Fig. S1C**). However, interpretation was limited due to 60% mortality in infected diabetic mice prior to this timepoint (**Fig. S1D**). Taken together, these findings indicate that *E. faecium* undergoes acute replication in control wounds but not in diabetic wounds, where it instead persists at higher levels at later timepoints of infection.

**Fig. 1:**
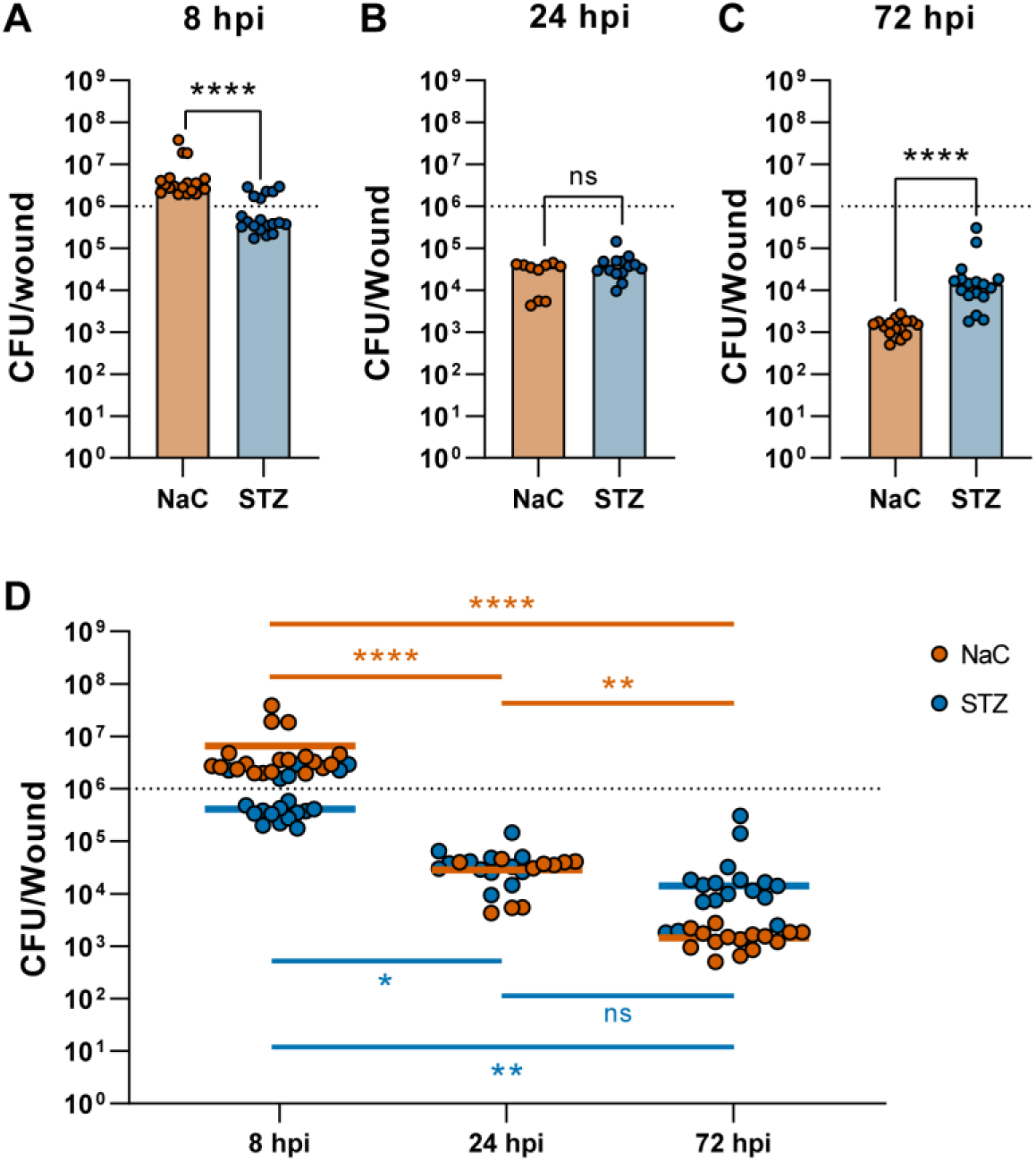
*E. faecium* persists at higher numbers at 72 hpi in diabetic mice. (**A-C**) CFU enumeration of control (NaC) and diabetic (STZ) mouse wounds infected with *E. faecium* (10^6^ CFU) at **(A)** 8 hpi, **(B)** 24 hpi and **(C)** 72 hpi. All CFU counts were enumerated on BHI + vancomycin (50 μg/ml). Bars represent median of n = 10-19 mice per group combined from 2-3 independent experiments. Statistical significance was determined using Mann-Whitney test. **** p<0.0001, ns: not significant. **(D)** Time course analysis of wound CFU counts from (A-C). Statistical significance between different timepoints of the same infection group was assessed with 2-way ANOVA mixed model analysis; Dotted line represents *E. faecium* inoculum (10^6^). * p<0.05, ** p<0.01, and **** p<0.0001, ns: non-significant.

### *E. faecium* co-infection facilitate the expansion of some co-infecting organisms

During CFU enumeration, we plated wound homogenates on both selective (BHI + 50 μg/mL vancomycin) and non-selective BHI agar. While CFU were comparable on both media at 24 hpi (**Fig. S2A**), by 72 hpi higher numbers were recovered on non-selective agar (**Fig. 2A**), suggesting the presence of a co-colonizing species. Plating on chromogenic agar confirmed the presence of both *Enterococcus* and *Staphylococcus* (**Fig. 2B**). The presence of *Staphylococcus* was unique to diabetic wounds at 72 h, regardless of infection status (**Fig. S2B**). 16s rRNA sequencing identified the co-colonizer to be *Staphylococcus lentus*, a murine skin commensal with pathogenic potential in animals that is rarely implicated in human infection [24–26]. Thus, diabetic wounds uniquely supported colonization by the commensal *S. lentus*.

**Fig. 2:**
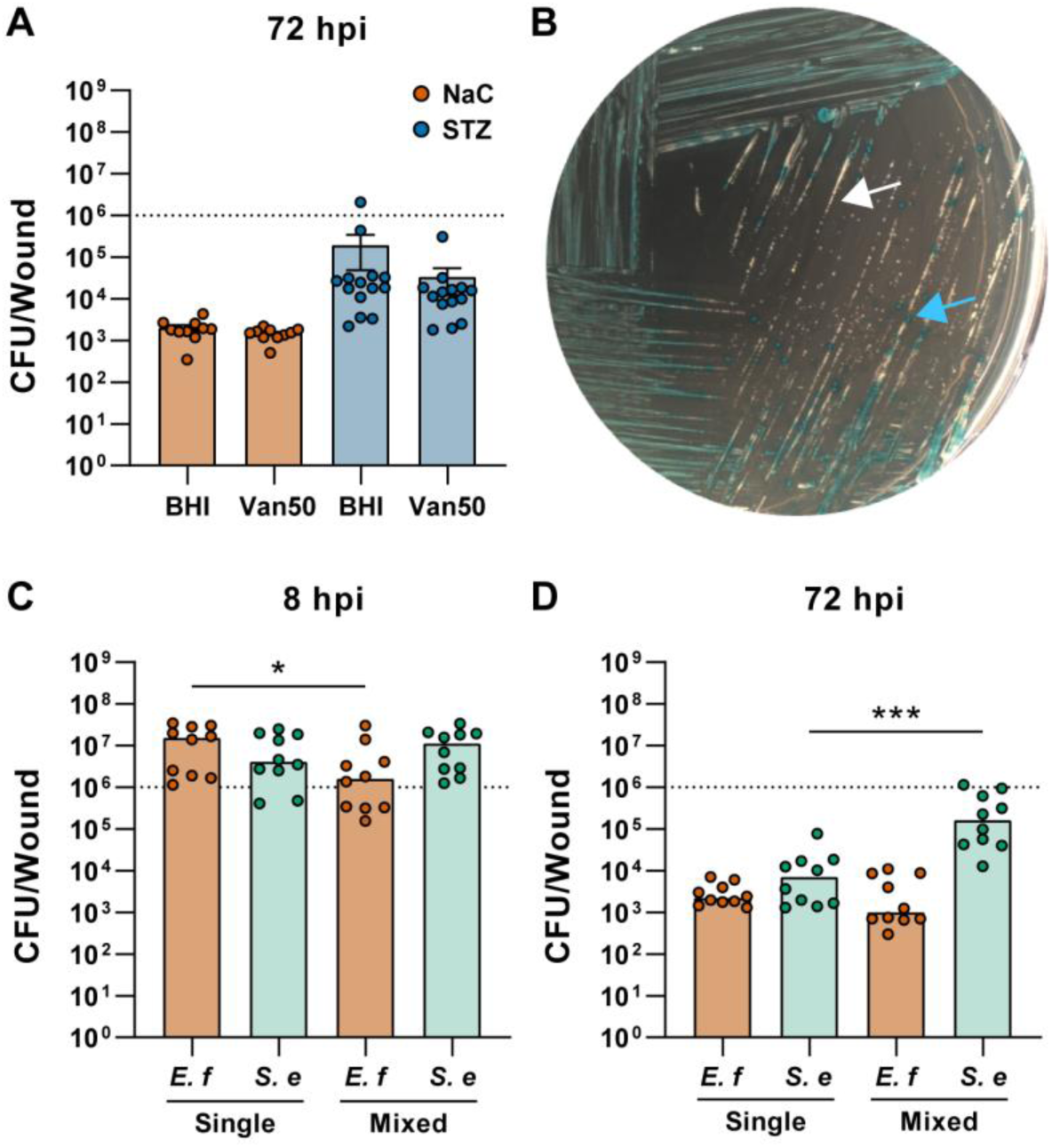
Mixed species wound infections enhance *S. epidermidis* growth. **(A)** 72 hpi wound CFU from control (NaC) and diabetic (STZ) wounds, plated on non-selective BHI and BHI + vancomycin (50 μg/ml) (Van50) agars. Bars represent mean ± SEM from n = 10-14 mice per infection group combined from 2 independent experiments. **(B)** UTI clarity streak plate from diabetic wound homogenates. Blue colonies (blue arrow) = *E. faecium*, white colonies (white arrow) = *S. lentus*. Plate image is representative of observations across 3 independent experiments. **(C-D)** Mixed species infection of control wounds with 1:1 *E. faecium* (*E.f)* and *S. epidermidis (S.e)* (10^6^ each). Wound CFU from **(C)** 8 hpi and **(D)** 72 hpi were quantified on BHI and Van50 agars. Bars represent median from n = 10 per infection group combined from 2 independent experiments. Statistical significance between BHI and vancomycin plate counts at each timepoint (A) or species-wise single and mixed species infection comparisons (C-D) was determined by Mann-Whitney Test. * p<0.05, ** p<0.01, *** p<0.001, **** p<0.0001, ns: non-significant.

To test whether *S. lentus* influenced *E. faecium* infection, we compared single and mixed infections in control mice. Although *S. lentus* CFU were significantly higher at 8 hpi in mixed species infection, there was no significant difference at 72 hpi when naturally occurring staphylococcal co-colonization was observed (**Fig. S2B-D**). By contrast, *E. faecium* CFU were significantly lower in mixed species infection at 8 hpi but comparable in both single and mixed species infections at 72 hpi. We next repeated the experiment with *Staphylococcus epidermidis*, a closely related human skin commensal and opportunistic pathogen [27, 28]. In this case, we observed similar *S. epidermidis* CFU at 8 hpi, while *E. faecium* CFU were significantly lower (**Fig 2C**). However, at 72 hpi *S. epidermidis* CFU were significantly higher in mixed compared to single species infection, whereas *E. faecium* CFU were comparable in both infections (**Fig. 2D**). These data suggest that *E. faecium* co-infection may facilitate the expansion of other opportunistic pathogens such as *S. epidermidis* in the wound environment.

### Sustained neutrophil, but not macrophage, infiltration in *E. faecium-*infected diabetic wounds

We hypothesized that differences in *E. faecium* clearance between diabetic and control mice might reflect differences in the innate immune response. To test this, we profiled wound infiltrates by flow cytometry. Single-cell suspensions were stained and gated into neutrophil (CD11b^+^ Ly6G^+^ F4/80^-^) and macrophage (CD11b^+^ Ly6G^-^F4/80^+^) populations **(Fig. S3**), and absolute cell numbers were quantified. At 24 h, we observed significantly more neutrophils in infected versus sterile diabetic wounds (**Fig. 3A**). We observed a similar trend in control mice, and overall neutrophil numbers were elevated in diabetic wounds in both conditions. At 72 h, although neutrophil infiltration was comparable in sterile and infected control mice, neutrophil infiltration was significantly higher in infected compared to sterile diabetic wounds, consistent with higher *E. faecium* CFU in diabetic wounds at this timepoint (**Fig. 1C**).

**Fig. 3:**
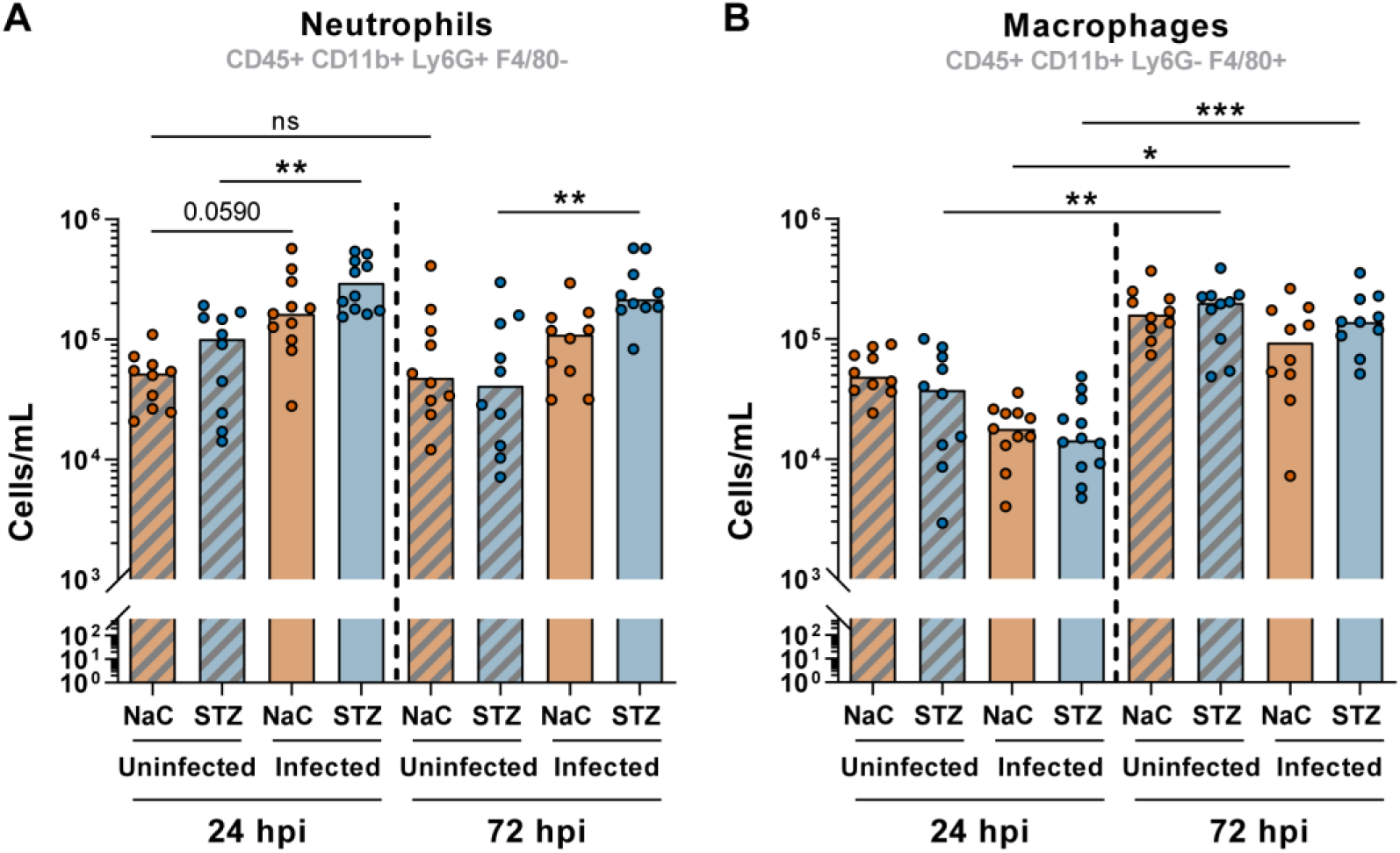
*E. faecium* infection elevates neutrophil infiltration in diabetic wounds, with macrophages elevated in 72 h wounds regardless of infection. Quantification of **(A)** neutrophil and **(B)** macrophage cell numbers from single cells dissociated from 24 and 72 hpi wounds. Bars indicated represent medians from n = 10 mice per infection group combined from 2 independent experiments. Statistical significance was determined by Kruskal Wallis test with Dunn’s multiple comparison test. p<0.05, ** p<0.01, *** p<0.001, ns: non-significant.

By contrast, macrophage numbers were comparable across all groups at 24 h (**Fig. 3B**). By 72 h, macrophage numbers had significantly increased relative to 24 h in both control and diabetic mice, independent of infection status (**Fig. 3B**), suggesting that wounding alone was sufficient to drive macrophage recruitment. Accordingly, CD45+ immune cell numbers, including neutrophils and macrophages, were consistently higher in wounded tissues compared to naïve cell skin (**Fig. S4**). Minor differences were noted in baseline immune cell levels between diabetic and control mice, but these were not statistically significant. Together, these findings show that while macrophage recruitment is largely wound-driven, *E. faecium* infection promotes sustained neutrophil infiltration in diabetic wounds.

### *E. faecium* wound infection induces transient pro-inflammatory cytokine production

We have previously shown that *E. faecalis* modulates host cytokine levels during wound infection [29], but the impact of *E. faecium*, particularly in a diabetic context, has not been investigated. We therefore measured pro-inflammatory cytokines and chemokines in wound lysates at 8 and 72 hpi (**Fig. 4, Fig. S5**). At 8 hpi, *E. faecium* induced robust TNF-α and IL-6, although levels were significantly lower in diabetic mice (**Fig. 4A-B**). IL-1α and IL-1β were similarly induced in response to infection, but IL-1α was significantly induced only in diabetic mice (**Fig. 4C-D**). G-CSF and GM-CSF were present at higher levels in response to infection at 8 h, although G-CSF levels were significantly lower in infected diabetic mice compared to control at the same timepoint (**Fig. 4E-F**). The anti-inflammatory cytokine IL-10 was comparably elevated in both control and diabetic groups in response to infection (**Fig. S5A**), whereas IL-17 and IFN-γ were present at lower levels in diabetic mice (**Fig. S5B-C**). Similarly, chemokines such MCP-1 and MIP-1α were also increased in response to infection at 8 h, and MCP-1 levels were significantly lower in diabetic wounds (**Fig. S5D-E**). By 72 hpi, cytokine levels were similar across all four experimental groups, with generally comparable or lower levels compared to 8 hpi timepoints particularly for infected groups (**Fig. 4, Fig. S5**). These findings show that while *E. faecium* induces a broad cytokine and chemokine response during acute infection, these responses are generally blunted in diabetic mice.

**Fig. 4:**
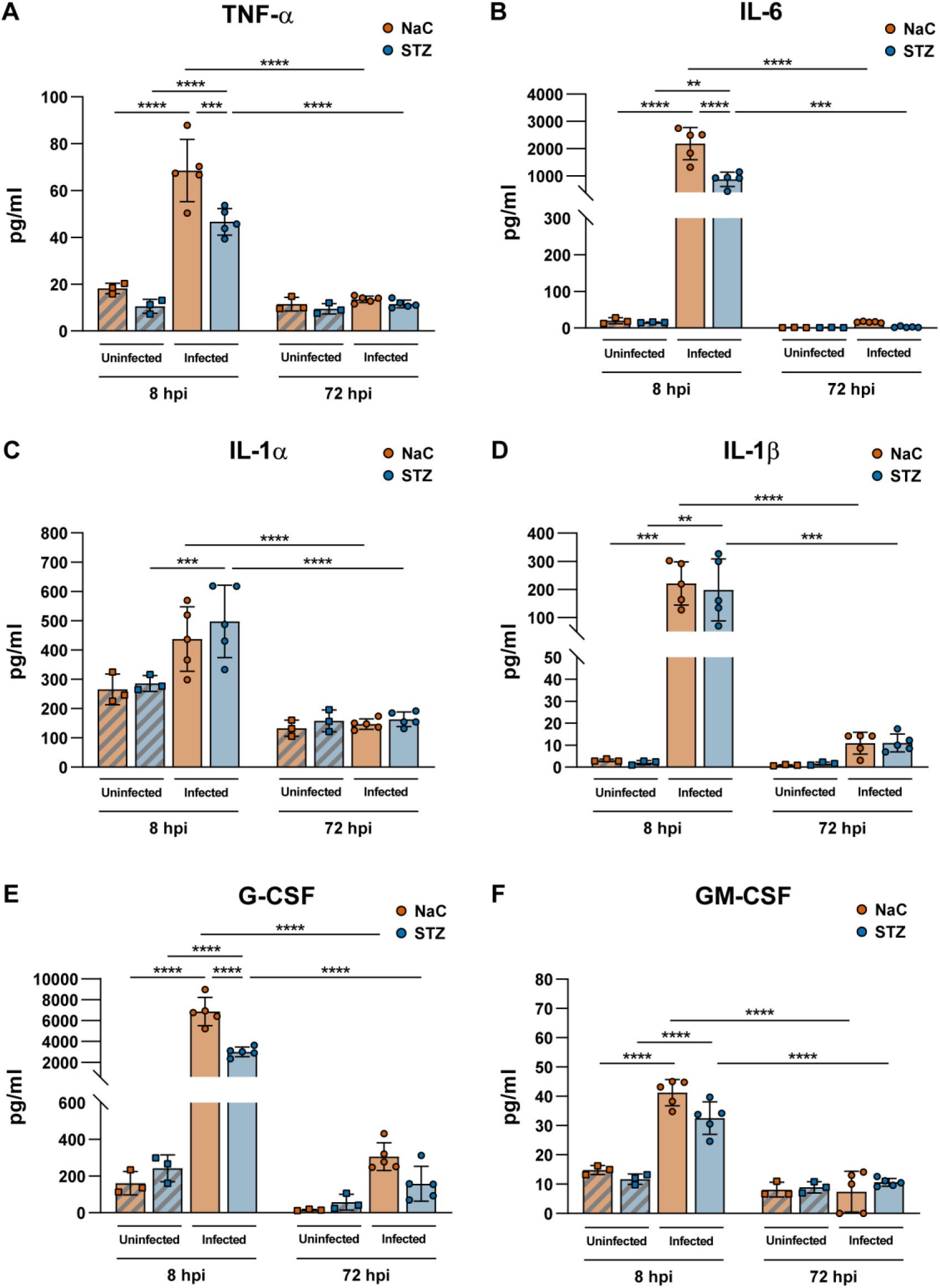
*E. faecium* wound infection triggers an acute inflammatory cytokine response that resolves by 72 h. Cytokine 23-plex assay of **(A)** TNF, **(B)** IL-6, **(C)** IL-1α, **(D)** IL-1β, **(E)** Granulocyte-Colony Stimulating Factor (G-CSF) and **(F)** Granulocyte, Monocyte-Colony Stimulating Factor (GM-CSF) in various wounds at 8 hpi and 72 hpi. Bars represent mean ± SD in n = 3-5 mice per infection group from one independent experiment. Statistical significance was determined by 2-way ANOVA with Tukey’s multiple comparisons test. * p<0.05, ** p<0.01, *** p<0.001 and **** p<0.0001, ns: non-significant. Only meaningful comparisons with p<0.05 are annotated.

### *E. faecium* infection delays wound healing in diabetic tissues

Macroscopic and histological examination of wounds can provide insights into wound healing during infection. *E. faecalis* can impede wound healing in non-diabetic mice, with epidermal hyperplasia or hyper-thickened epidermis serving as a marker of impaired healing [30]. To assess whether *E. faecium* impacts healing in a diabetic context, we performed hematoxylin and eosin (H&E) staining and macroscopic examination at 72 h across four groups: uninfected control and diabetic wounds (**Fig. 5A-B**) and *E. faecium* infected control and diabetic wounds (**Fig. 5C-D**). Wounds from uninfected control (**Fig. 5A**) and diabetic (**Fig. 5B**) mice exhibited re-epithelization, with new epidermis advancing from the wound edges (black arrowheads, **Fig. 5A-B)**. Uninfected diabetic wounds also showed a hyper-thickened epidermis, indicating impaired healing even in the absence of infection (yellow double arrow, **Fig. 5B**). In *E. faecium*-infected wounds, control mice similarly exhibited a hyper-thickened epidermidis compared with adjacent old epidermis (yellow double arrows, **Fig. 5C-D**). Furthermore, dense infiltrates of small, round nucleated cells beneath the wound bed (red arrows) suggest actively infiltrating immune cells. Diabetic infected wounds exhibited equally pronounced hyper-thickening and immune infiltration (**Fig. 5D**). Thus, both the diabetic environment and *E. faecium* infection appear to impact the progression of wound healing. These findings demonstrate that both diabetes and *E. faecium* infection independently impair wound healing, with overlapping pathological features including epidermal hyper-thickening and excessive immune infiltration.

**Fig. 5:**
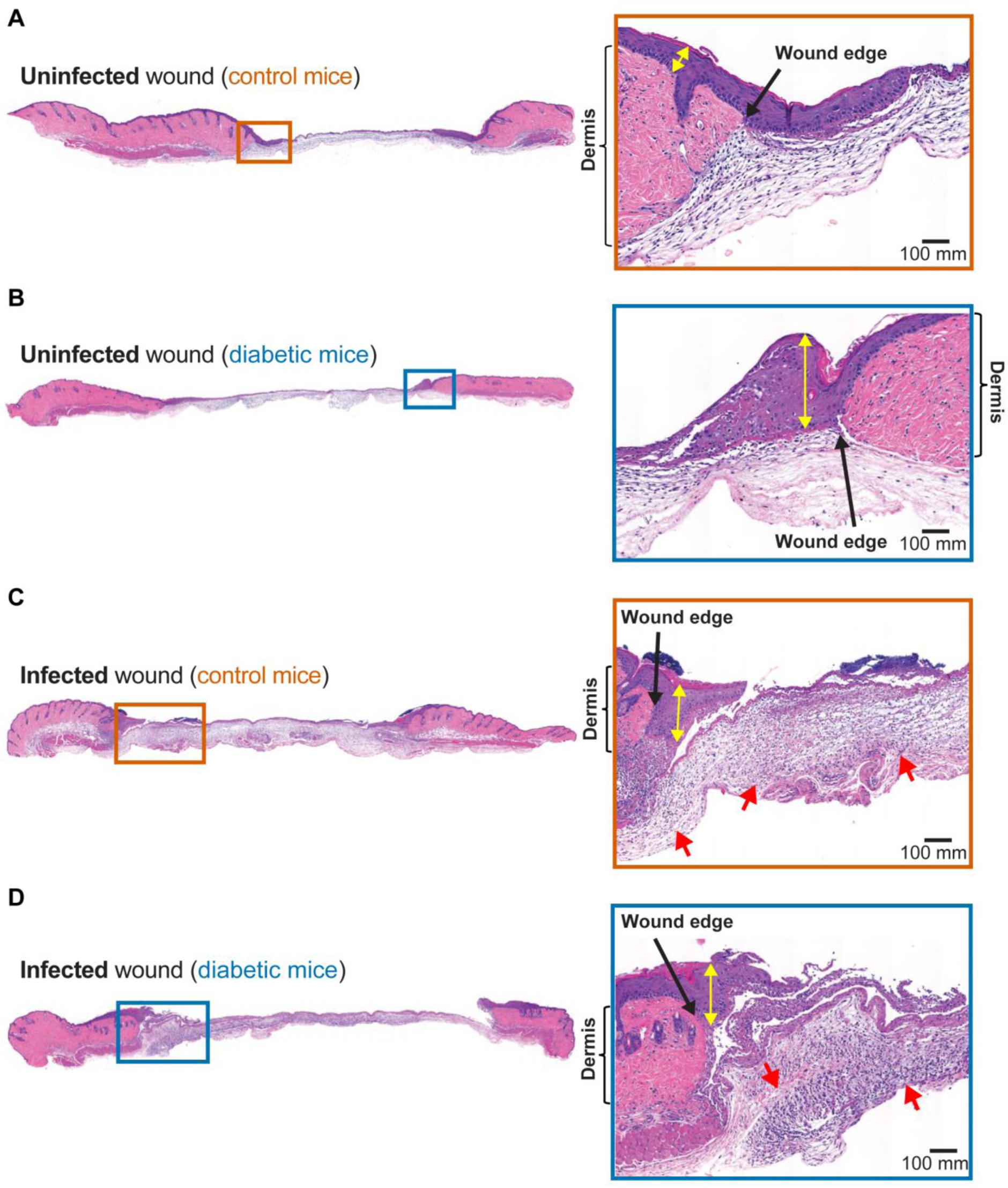
*E. faecium* infected and diabetic wounds exhibit epithelial hyper-thickening with evidence of immune cell infiltration during infection. Hematoxylin and Eosin staining of 5 mm sections from **(A)** Uninfected-control, **(B)** Uninfected-diabetic, **(C)** Infected-control and **(D)** Infected-diabetic wounds at 72 hpi. Right insets show magnified view around the respective wound edges (black arrows). Yellow double arrows: epithelial thickness. Red arrows: regions of immune cell infiltration.

## Discussion

The global rise in diabetes, coupled with the emergence of multidrug-resistant *E. faecium,* underscores an urgent need to understand the dynamics of *E. faecium* wound infection in diabetic populations [8, 11, 14, 31]. Here we show that diabetic wounds do not support acute replication of *E. faecium* after inoculation. Instead, bacterial colonization stabilizes at inoculum levels, while control mice exhibit rapid bacterial replication followed by efficient clearance. These findings suggest that diabetes-associated changes in the wound environment, potentially involving immune dysfunction or altered skin microbiota, constrain early bacterial expansion but permit longer-term persistence.

Although diabetes is classically associated with a pro-inflammatory baseline, marked by elevated plasma cytokines [32], our data reveal that the initial cytokine response to wound infection is blunted in diabetic mice. Despite this dampened cytokine response, *E. faecium* persists in diabetic wounds alongside sustained neutrophil infiltration, which is not mirrored by elevated cytokine levels or increased macrophage recruitment at later timepoints of infection. The coexistence of persistent *E. faecium* CFU and the sustained presence of neutrophils suggest a possible functional impairment of neutrophils in diabetic wounds, consistent with literature describing altered NETosis and phagocyte dysfunction in diabetes [33–35].

Microbial factors can further drive immune dysregulation. While not well-studied for *E. faecium, E. faecalis*, which is also commonly associated with human disease, can suppress elements of innate immunity. For instance, *E. faecalis* persists in wounds via a combination of reduced NF-κB signaling in macrophages, reduced phagocyte degranulation or phagolysosome formation, and even replication within macrophages and neutrophils [29, 36–40]. *E. faecalis* also impacts wound healing via impairment of epithelial to mesenchymal transition, a process essential for wound remodeling and re-epithelialization, even in non-diabetic wounds, providing precedence for direct modulation of wound healing mechanisms [41]. Additionally, *E. faecalis*-derived reactive oxygen species (ROS) has been linked to impaired wound healing [42]. These immune and tissue-modulatory effects likely extend to *E. faecium*, permitting both it and possible opportunistic partners to persist in the otherwise hostile wound microenvironment.

Polymicrobial interactions are a defining feature of chronic wounds, particularly in diabetes [19, 43, 44]. This study identified commensal *S. lentus* as a co-colonizer exclusive to diabetic wounds, suggesting that diabetic host microenvironmental changes may favor colonization by commensal or opportunistic skin species. While *S. lentus* did not impact *E. faecium* persistence, mixed infections with *S. epidermidis,* a human skin commensal, led to augmented growth of the latter, suggesting that *E. faecium* may facilitate opportunist expansion through niche modification or immune modulation. This finding parallel reports increased staphylococcal density and reduced overall diversity in diabetic mice skin [45–48]. Analogous polymicrobial occurrences were reported in group B streptococcal diabetic wound infections, where *E. faecalis* and *Staphylococcus xylosus* were recovered from wound homogenates, underscoring both the importance using non-selective media when isolating bacteria from diabetic hosts and the need to consider polymicrobial synergy in these infection settings [49]. Although there is currently no evidence that *E. faecium* can modulate the growth of co-infecting species during wound infection, there is precedence from studies in which *E. faecalis* which augments *E. coli* CFU in wounds [50] or *E. faecalis* growth is enhanced by *Staphylococcus aureus* [51].

Further supporting the multifactorial nature of chronic wound infection, our histological data revealed hyper-thickened epidermis and impaired fibroblast infiltration in both diabetic and control infected wounds. Diabetic wounds, on the other hand, display hyper-thickened epidermis regardless of the infection status, indicating the diabetic environment may be the primary driver for delayed wound healing [52, 53]. This reinforces the concept that diabetes and bacterial infection independently—and likely synergistically—impair core regenerative processes, an outcome exacerbated in polymicrobial contexts by biofilm formation and cooperative immune evasion strategies.

In conclusion, this work reveals distinct host-pathogen dynamics in diabetic wound infections, where *E. faecium* persists despite robust initial immune responses, contributing to delayed wound healing independent of host diabetic status. Further work should dissect the molecular mechanisms underlying immune modulation and interspecies interactions to inform targeted therapeutic strategies for chronic diabetic wound infections colonized by *E. faecium*.

## Material and Methods

### Bacterial-strains and growth conditions

Vancomycin-resistant *Enterococcus faecium* strain E745 was grown overnight at 37°C in brain heart infusion broth (BHI; Neogen, Lansing, Michigan) without antibiotics. For mixed species experiments, *Staphylococcus epidermidis* strain ATCC 12228 and *Staphylococcus lentus* isolates from diabetic wound infections were grown overnight at 37°C in BHI broth without antibiotics. For wound infection experiments, overnight cultures of bacteria were washed and resuspended in PBS (Invitrogen, USA) to OD 0.4 (equivalent to 1×10^8^ CFU/ml) for infection.

### Streptozocin-induced diabetic mice model

Diabetes was induced in male C57BL/6J mice (5-6 weeks old) via intraperitoneal injection of freshly prepared streptozocin (STZ; Sigma, USA, Cat# S0130-50MG), administered at 50 mg/kg in 0.1 M sodium citrate solution (NaC; Sigma, USA, Cat# C7254-1KG), pH 4.5 daily for five consecutive days [23]. Control mice received only equivalent NaC solution, pH 4.5. Mice were fasted 4-5 hours prior to injection and left on a regular diet for seven days post-injection. Diabetes was confirmed 7 days post-injection by fasting blood glucose measurements (>300 mg/dl) using an ACCU-CHEK Instant Glucometer (Model 961; Roche International).

### Murine excisional wound infection model

Excisional wounds (6 mm) were created on the dorsal skin of anaesthetized 7-8 week-old C57BL/6J male mice (isoflurane), as previously described [30]. Hair was removed by shaving and depilatory cream, and skin was disinfected with ethanol, prior to wounding. Wounds were inoculated with 10^6^ cells of *E. faecium* (E745) either alone or in combination (1:1) with *S. epidermidis* or *S. lentus* (10^6^ cells/species). Tegaderm dressing (3M, St Paul Minnesota, USA) was applied, and mice were monitored regularly prior to euthanization at 8, 24 or 72 hpi.

### CFU enumeration from wound homogenates

Excised wounds (including Tegaderm) were homogenized in 1 ml PBS using a Lysing Matrix M (MP Biomedicals, USA, Cat# 6923050) at 4.0 m/s for 20 seconds for five rounds. Wound homogenates were serially diluted in PBS, and serial dilutions were plated on BHI agar with or without 50 μg/ml of vancomycin (Van50) (Merck, USA, Cat# 1709007) for CFU enumeration. The vancomycin-resistant *E. faecium* CFU counts were taken from Van50 plates alone, while vancomycin-susceptible *S. epidermidis* and *S. lentus* CFU counts were taken as the difference between total BHI counts and *E. faecium*-specific Van50 counts.

### Cytokine analysis

After homogenization as above, wound lysates were diluted 1:2 and analyzed for total cytokine concentrations using the Bio-Plex PRO Mouse Cytokine 23-plex Assay (Bio-Rad, California, USA), following manufacturer’s instructions as previously described [54].

### Histology

Excised wound tissues (Tegaderm removed) were fixed in 4% paraformaldehyde (PFA; Sigma, USA) for 24 h at 4°C. Fixed tissues were embedded in paraffin, sectioned at 5 mm thickness, stained with hematoxylin and eosin (H&E), and imaged with the help of Advanced Molecular Pathology Laboratory at the Institute of Molecular and Cell Biology, A*STAR, Singapore. Slide scanned images were analyzed via the Carl Zeiss Zen Lite software (Carl Zeiss, Germany).

### Flow cytometry

Excised wounds tissues were digested with 0.2 mg/ml Liberase TL (Merck, USA) at 37°C for 1 h, and single-cell suspensions filtered through a 40 μm cell strainer (SPL Life Sciences, South Korea). The single cell suspension was normalized to a cell count of 1×10^7^ cells/ml. 50 μl of the suspension was blocked with 1:100 TruStain FcX PLUS anti-mouse CD16/32 (BioLegend, USA, Cat# 156604) antibodies for 30 min on ice and then immediately stained for 1 h on ice with the following antibodies, each at 1:100 dilution: BV510 anti-mouse CD45 (BioLegend, Cat# 103138, USA), FITC anti-mouse Ly6G (BioLegend, Cat# 127606, USA), APC anti-mouse F4/80 (BioLegend, Cat# 123116, USA), PE anti-mouse/human CD11b (BioLegend, Cat# 101208, USA). After fixation with 4% PFA, samples were analyzed using a 5-laser BD LSRFortessa X-20.

For absolute counts of immune cell infiltrates, AccuCheck Counting Beads (Invitrogen, Cat# PCB100, USA) were added immediately before flow cytometry in a ratio of 100 μl beads to 400 μl of single-cell suspensions, according to manufacturer protocols. Absolute counts were calculated by normalizing the total number of cells to the total number of beads acquired (Fig S3).

### Data and statistical analysis

Flow cytometry data were plotted and analyzed using FlowJo v10.8.0 (Becton Dickinson, USA). Other graphs were created and statistical analyses performed with GraphPad Prism (Version 10.01 for Macintosh OS). Details on specific tests applied and data obtained are contained within figure legends.

### Ethics declaration

All procedures were approved and performed in accordance to the requirements of Institutional Animal Care and Use Committee in Nanyang Technological University (ARF SBS/ NIEA0198Z). Diabetic wound infections were performed in accordance to approved animal use protocol number A19061/A24066.

## Acknowledgements

This work was conducted at the Singapore Centre for Environmental and Life Science Engineering (SCELSE), whose research is supported by the National Research Foundation Singapore, Ministry of Education, to Nanyang Technological University and the National University of Singapore under its Research Centre of Excellence program. This work was funded by a Singapore Ministry of Education Academic Research Fund Tier 2 Grant awarded to K.A.K. (MOE2019-T2-2-089).

We would like to thank Willem van Schaik and Ross McInnes for providing the E745 strain, and for insightful discussions about this project. We would also like to thank the Advanced Molecular Pathology Lab at A*STAR, Singapore for their assistance in processing the histology slides, and Bio-Rad service staff for their assistance in running the Cyto-Plex measurements. We are grateful to Julien Huttman for critical reading of the manuscript

## Author contributions

N.J designed and performed the experiments and wrote the initial draft of the manuscript. F.R.T helped in the design and analysis of flow cytometry experiments, and provided figure visualizations as well as edited the manuscript. H.A helped in the design of the animal infection model and experiments, and edited the manuscript C.N.J.Y and R.T.J.W assisted in animal model experiments. D.L.B provided expertise in the analysis of wound histology. C.J.S helped in the design, performance and analysis of experiments and editing the manuscript. K.A.K conceptualized the project, obtained funding, managed the project and edited the manuscript. All authors read and agreed on the published version of the manuscript.

---

**Supplementary Fig. 1:**
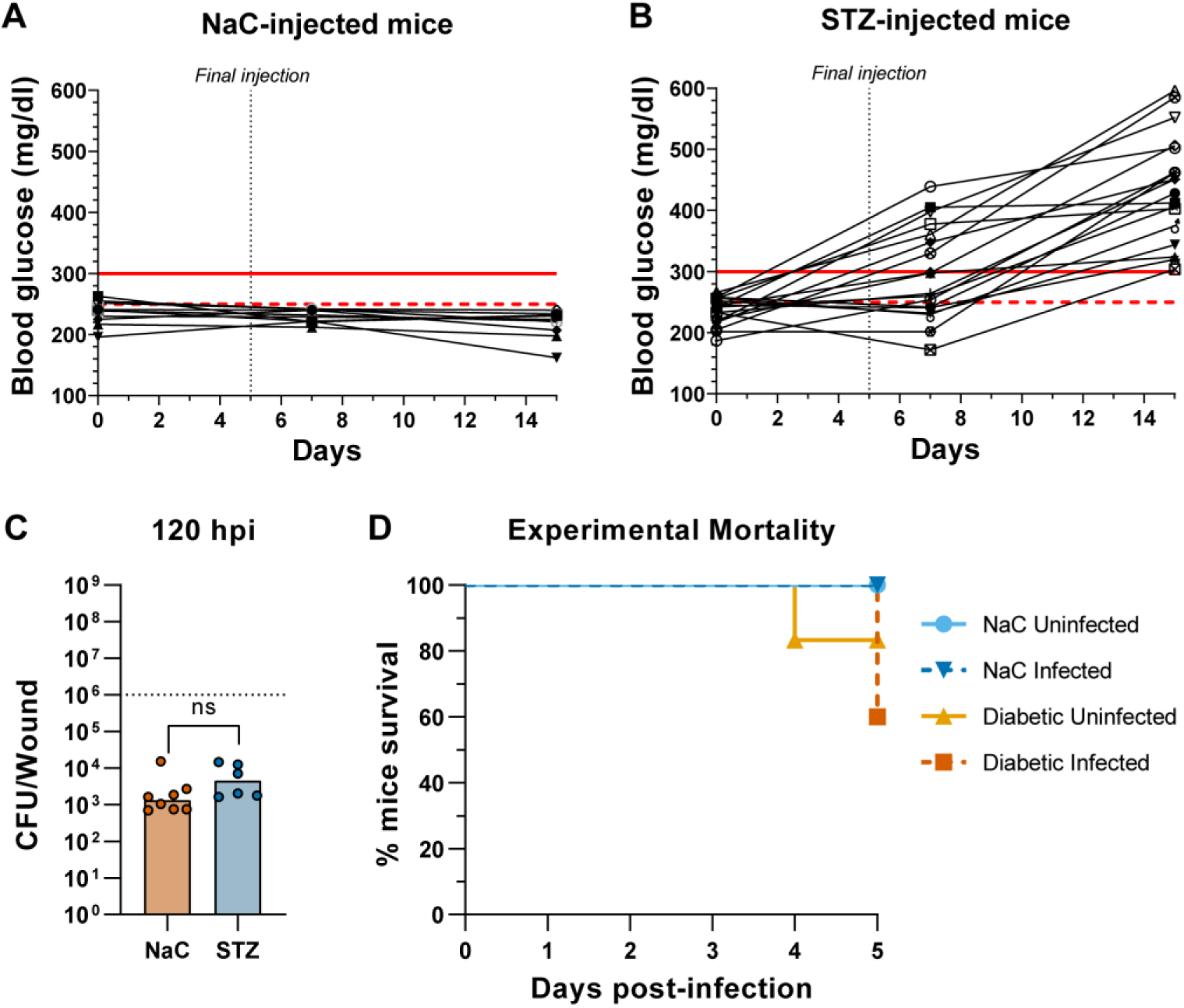
STZ treatment induces hyperglycemia and increases mortality following wounding and infection. **(A-B)** Fasted mice were injected with either **(A)** sodium citrate (NaC) or **(B)** streptozocin (STZ) for 5 days, and monitored for fasting blood glucose concentrations at 0, 7 and 15 days. Each data point depicts matched blood glucose measurements from n = 18 mice combined from 3 independent experiments. Red dashed and solid lines indicate hyperglycemic (>250 mg/dl) and diabetic (>300 mg/dl) thresholds respectively. **(C)** Wound CFU of control (NaC) and diabetic (STZ) mice at 5 dpi on BHI + vancomycin (50 μg/ml). Bars represent mean ± SD from n = 6-8 per infection group from one independent experiment. Dotted line represents *E. faecium* inoculum (10^6^). Statistical significance was determined by Mann-Whitney Test. ns = non-significant. **(D)** Mortality of mice up to 5 dpi from one independent experiment (n = 10 per mice group), showing percentage survival of control (NaC) mice or diabetic mice with or without *E. faecium* infection.

**Supplementary Fig 2:**
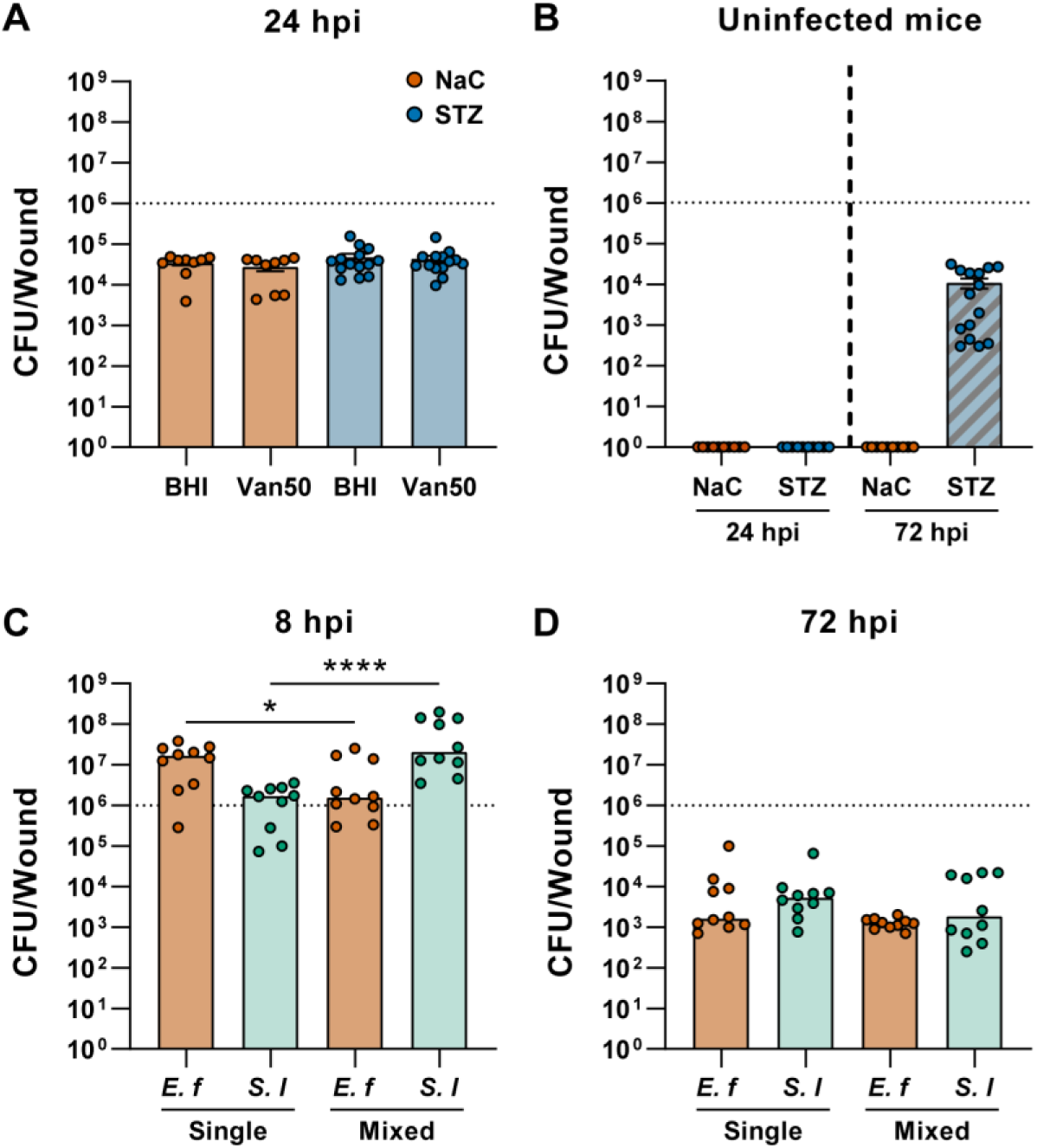
*S. lentus* uniquely colonizes diabetic wounds at 72 hpi without altering mixed species growth dynamics. All wound homogenates were plated on plated on non-selective BHI and BHI + vancomycin (50 μg/ml) (Van50) agars. **(A-B)** Wound CFU from control (NaC) and diabetic (STZ) wounds after **(A)** 24 h of infection with *E. faecium*, or **(B**) 24 or 72 h of mock-infection with PBS. Bars represent median from n = 10-15 mice per infection group combined from 2-3 independent experiments. **(C-D)** Mixed species infection of NaC wounds with 1:1 *E. faecium* (*E.f)* and *S. lentus (S.l)* (10^6^ each). Wound CFU from **(C)** 8 hpi and **(D)** 72 hpi were quantified on BHI and Van50 agars. Bars represent median from n = 10 per infection group across 2 independent experiments. Statistical significance between single and mixed species infections was determined by Mann-Whitney Test. * p<0.05, **** p<0.0001.

**Supplementary Fig 3:**
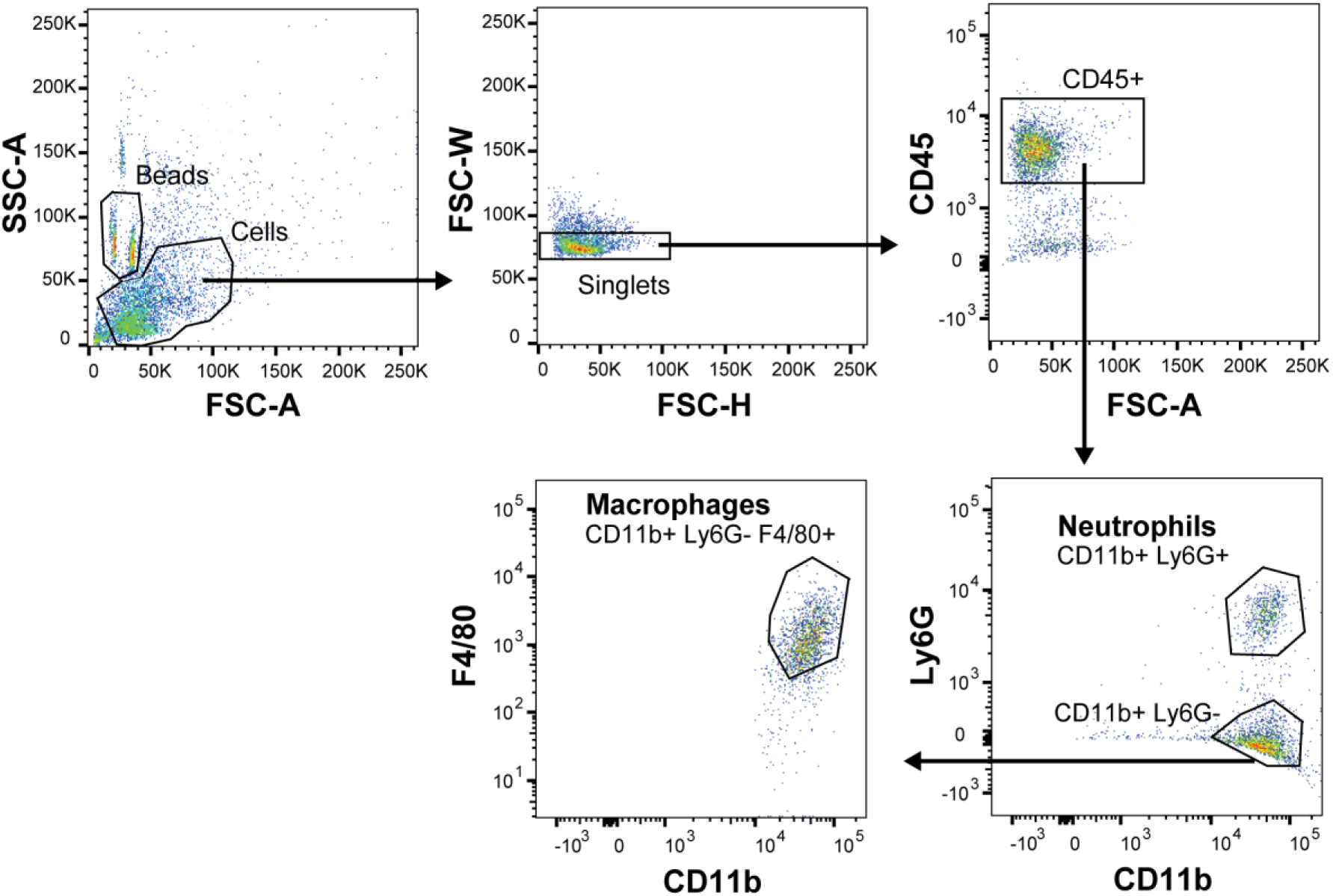
Flow cytometry gating strategy for quantifying immune cell infiltrate in wound homogenates. Single wound cells were gated from the FSC-A/SSC-A and FSC-A/FSC-W plots. Leukocytes or immune cells were then gated as the CD45^+^ population, from which Ly6G^+^CD11b^+^ (neutrophils) population is gated. The Ly6G^-^ cells were further gated to F4/80^+^CD11b^+^ (macrophages) population. To normalize quantification of wound cells, AccuCheck counting beads were gated and quantified from the FSC-A/SSC-A plot as indicated.

**Supplementary Fig 4:**
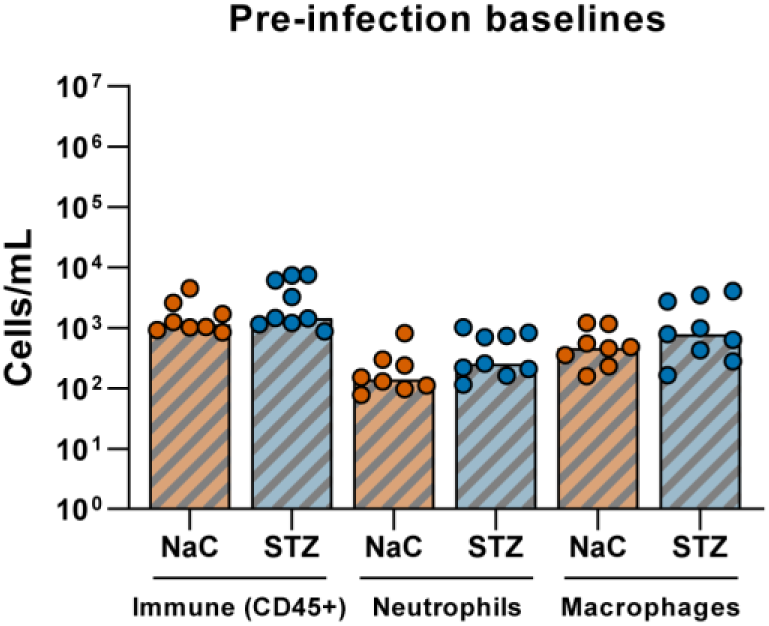
Baseline skin immune-cell populations do not differ between control and diabetic mice. Total numbers of immune cells (CD45^+^), neutrophils (CD45^+^CD11b^+^Ly6G^+^) and macrophages (CD45^+^Ly6G^-^CD11b^+^F4/80^+^) from control and diabetic mice skin prior to wounding and infection. Bars represent median from n = 8-9 mice per infection group combined from 2 independent experiments. Statistical significance was determined by Kruskal Wallis Test with Dunn’s multiple comparisons test. All comparisons were found to be non-significant (ns).

**Supplementary Fig 5:**
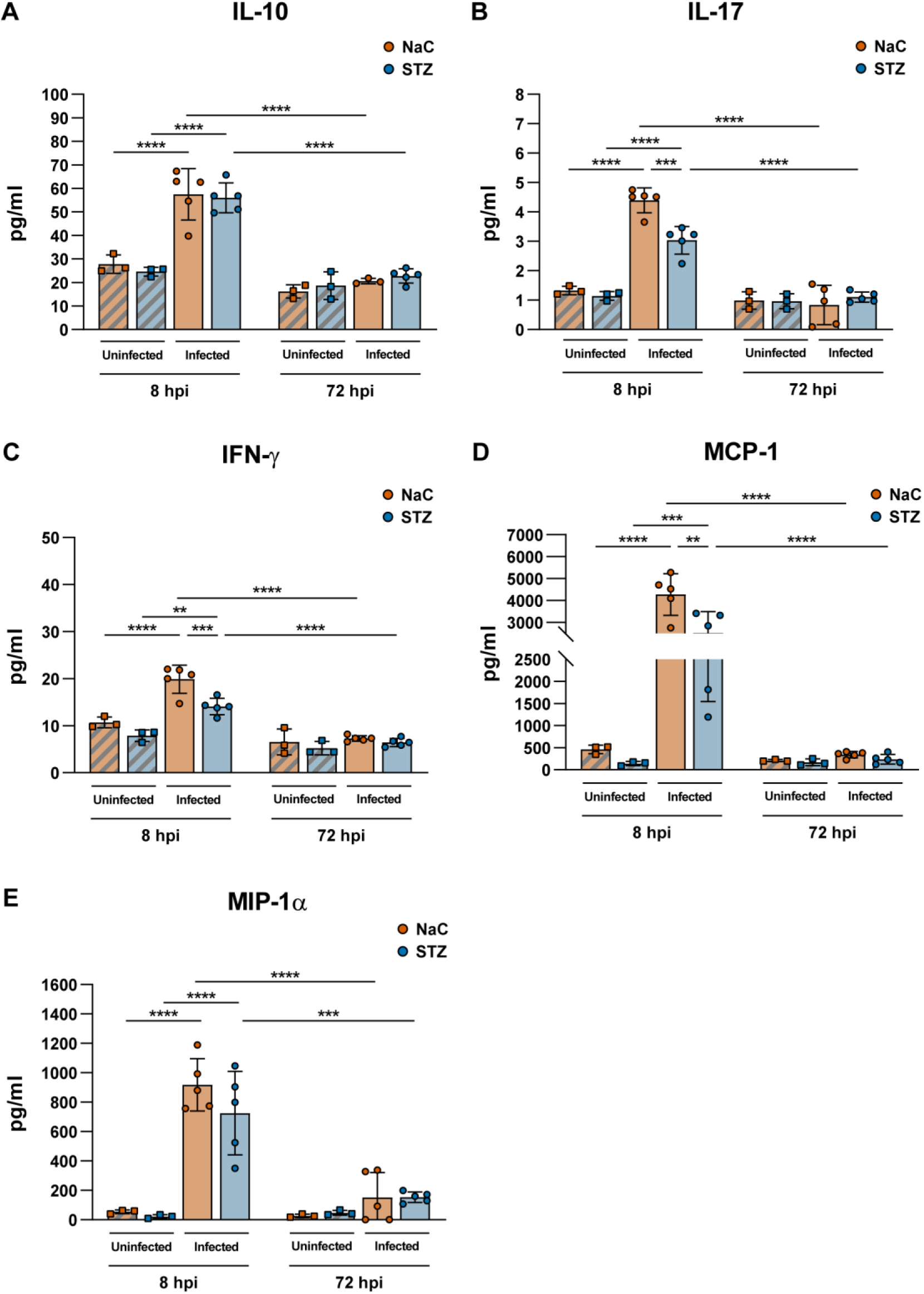
Cytokine levels peak at 24 hours following *E. faecium* infection and diminish by 72 hours. Cytokine 23-plex assay of **(A)** IL -10, **(B)** IL-17, **(C)** IFN **(D)** MCP-1, **(E)** MIP-1α in various wounds at 8 hpi and 72 hpi. Bars represent mean ± SD from n = 3-5 per infection group from one independent experiment. Statistical significance was determined by 2-way ANOVA with Tukey’s multiple comparisons test. * p<0.05, ** p<0.01, *** p<0.001 and **** p<0.0001, ns; non-significant.

